# Bromodomain protein IBD1 bridges histone acetylation and H2A.Z deposition to fine-tune transcription and prevent hyperactivation in *Tetrahymena thermophila*

**DOI:** 10.1101/2025.06.29.662184

**Authors:** Zhe Zhang, Haicheng Li, Fan Wei, Aili Ju, Hongzhen Jiang, Fei Ye, Yongqiang Liu, Shan Gao, Yuanyuan Wang

## Abstract

Eukaryotic gene expression is dynamically regulated through the interplay between histone modifications and chromatin remodeling, yet how these processes are coordinated remains incompletely understood. Here, we identify IBD1 as a critical adaptor that bridges histone acetylation and SWR-mediated H2A.Z deposition. We demonstrate that the bromodomain of IBD1 recognizes histone acetylation, specifically H3K9/14 di-acetylation to recruit the SWR complex subunit ARP6, ensuring precise H2A.Z incorporation into chromatin. Genetic disruption of IBD1, either by deletion or bromodomain mutation, attenuates H2A.Z occupancy at target loci, confirming its essential role in H2A.Z deposition. Strikingly, perturbation of the histone acetylation-IBD1-H2A.Z axis triggers transcriptional hyperactivation, revealing a dual function of H2A.Z in sustaining basal transcription while preventing aberrant gene induction. This defines a novel regulatory paradigm in which IBD1 couples acetyl-mark decoding to SWR-dependent H2A.Z deposition, establishing transcriptional homeostasis. Our findings resolve a central ambiguity by demonstrating that H2A.Z acts as a repressive buffer in highly acetylated regions, counterbalancing the inherent activation potential of acetylation. This study provides a framework for investigating H2A.Z functions and reveals new perspectives on how chromatin modifications and histone variants interplay in transcriptional regulation.

## Introduction

Gene transcription in eukaryotes is dynamically regulated by chromatin architecture, with nucleosomes acting as the fundamental units of compaction and signaling platforms (Luger et al 1997). Each nucleosome comprises an octamer of core histones (H2A, H2B, H3, H4) wrapped by ∼147 bp of DNA, and its regulatory potential is amplified by histone variants and post-translational modifications (PTMs) (Kouzarides 2007, Strahl & Allis 2000, Wong & Tremethick 2025). Among these, the H2A.Z variant, highly conserved from yeast to humans, is preferentially deposited near promoters and transcription start sites (TSSs), yet its transcriptional role remains context-dependent, mediating both activation and repression (Giaimo et al 2019, Kreienbaum et al 2022).

The ATP-dependent SWI2/SNF2-related (SWR) complex orchestrates H2A.Z deposition into chromatin, replacing canonical H2A-H2B dimers (Billon & Côté 2012, Brahma et al 2017, Obri et al 2014). Recent structural work has revealed that Yaf9, ENL, AF9, Taf14, and Sas5 (YEATS) domain of glioma amplified sequence 41 (GAS41), a subunit of mammalian SWR complexes, binds H3K14ac or H3K27ac to facilitate H2A.Z incorporation and transcriptional activation (Kikuchi et al 2023). Beyond YEATS domains, acetyl-lysine “readers” like bromodomains (BRDs) similarly bridge PTM decoding to chromatin remodeling. For instance, bromodomain and extra-terminal (BET) family proteins leverage BRD-ET domain synergies to recruit transcriptional machinery (Fujisawa & Filippakopoulos 2017, Zhang et al 2025). In yeast, Bdf1, a BET family member and the BRD-containing subunit of the SWR1 complex, targets the remodeler to acetylated H2A/H4 and interacts with the general transcription factor TFIID, thereby functionally bridging H2A.Z deposition with transcriptional activation (Altaf et al 2010, Matangkasombut et al 2000). However, whether H2A.Z deposition at acetylated loci amplifies or attenuates transcription remains unclear, hinting at uncharacterized feedback mechanisms.

The unicellular model organism *Tetrahymena thermophila* (hereafter referred to as *Tetrahymena*) provides a unique system to dissect these dynamics. Its transcriptionally active macronucleus (MAC) is enriched for H2A.Z and histone acetylation, while the silent micronucleus (MIC) lacks both marks (Allis et al 1980, Allis et al 1986, Wahab et al 2020). Prior studies in *Tetrahymena* implicated H2A.Z in coordinating transcription-associated modifications (e.g., H3K4me3) and DNA methylation analogues (6mA) (Wang et al 2025, Wang et al 2019). Despite these advances, how H2A.Z deposition fine-tunes transcriptional output is unresolved.

Here, we identify IBD1 as a critical adaptor that bridges histone acetylation and SWR-mediated H2A.Z deposition. We demonstrate that IBD1 recognizes histone acetylation, specifically H3K9/14 di-acetylation, to guide the H2A.Z deposition on chromatin by the SWR complex subunit ARP6. Disrupting IBD1, either through deletion or BRD mutation, impairs H2A.Z occupancy. Strikingly, perturbation of the histone acetylation-IBD1-H2A.Z axis leads to transcriptional hyperactivation, revealing the role of H2A.Z in maintaining basal transcription while preventing hyperactivation. Our findings establish a novel regulatory axis in which IBD1 links histone acetylation to SWR-dependent H2A.Z deposition, ensuring transcriptional homeostasis. This study resolves a key ambiguity regarding H2A.Z’s function by demonstrating its repressive role in highly acetylated regions. By elucidating the crosstalk between histone modifications and chromatin remodeling, our work advances the understanding of epigenetic mechanisms fine-tuning gene expression and provides a framework for future studies on transcriptional regulation.

## Results

### H2A.Z enrichment correlates with transcriptionally active genes

To investigate the functional role of H2A.Z in *Tetrahymena* (Table S1), we systematically analyzed its cellular localization. Using previously constructed N-terminally hemagglutinin (HA)-tagged H2A.Z (H2A.Z-NHA) cells (Wang et al 2017), immunofluorescence (IF) staining revealed that H2A.Z exhibited strong nuclear compartmentalization, being highly enriched in the transcriptionally active MAC while completely excluded from the transcriptionally silent MIC in vegetative stages (Fig. 1A), consistent with a potential role in transcriptional regulation.

**Fig. 1.**
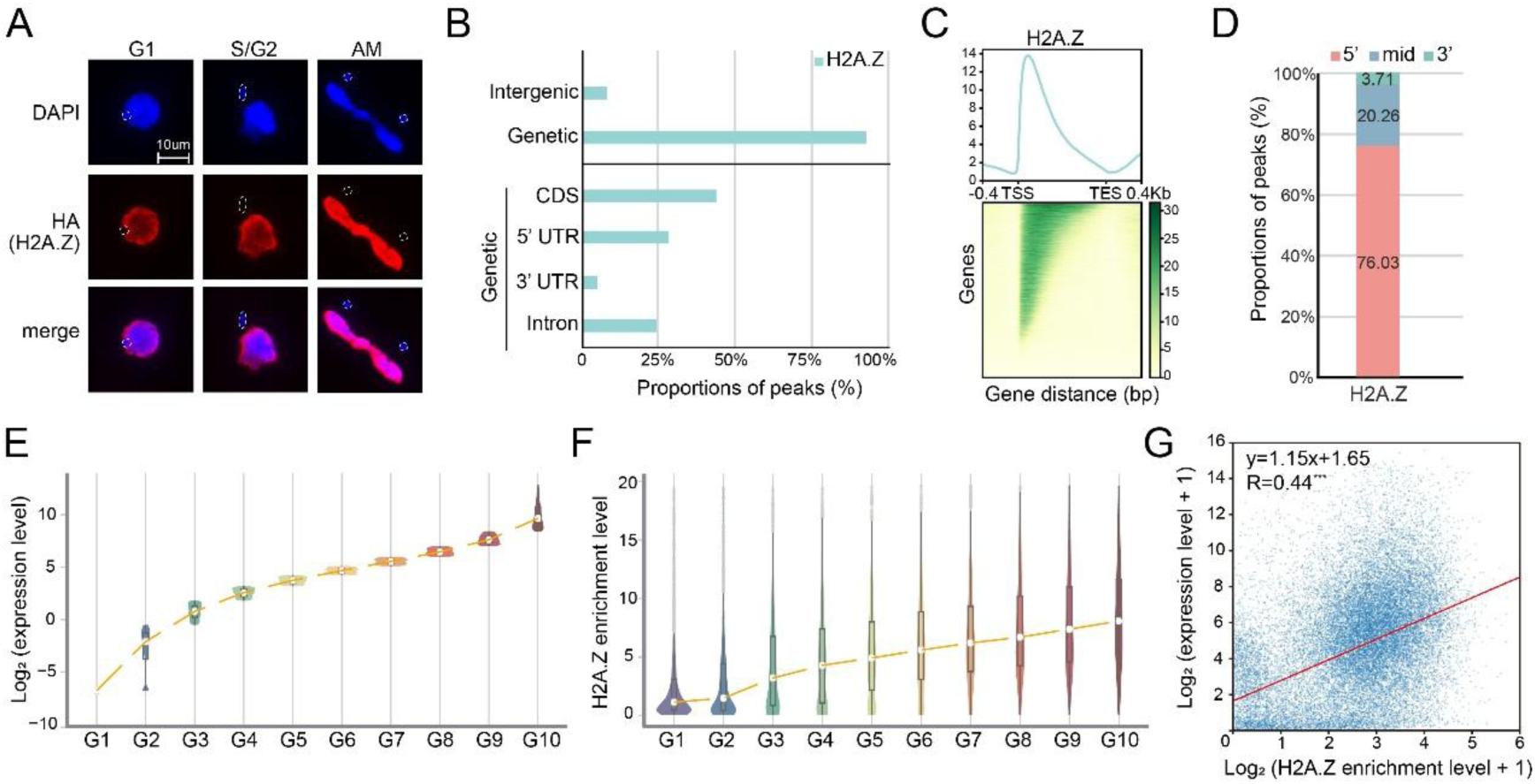
H2A.Z enrichment correlates with transcriptionally active genes. A. IF staining shows that HA-tagged H2A.Z proteins are exclusively localized to the macronucleus (MAC) but absent in the micronucleus (MIC), outlined with dotted circles. Nuclei were counterstained with DAPI (blue). B. Distribution of H2A.Z in the different genomic regions in wild-type (WT) cells indicates that it was predominantly enriched in genetic regions, especially coding sequences (CDS), compared to untranslated regions (UTRs) in the *Tetrahymena* genome. C. Distribution of H2A.Z along the gene body shows that it was preferentially enriched at the 5’ end of gene bodies in WT cells. Genes are scaled to unit length and extended to each side for 0.4kb. TSS, transcription start site. TES, transcription end site. D. Proportions of H2A.Z peaks distributed across different gene body regions. Genes were evenly divided into three segments: the 5′ part, the 3′ part, and the middle region (mid). The proportion of H2A.Z peaks in each segment relative to the total number of peaks was then calculated. E. Gene expression levels across ten groups in WT cells. y axis: Log_2_-transformed expression levels. x axis: 10 quantiles of genes rank by their gene expression levels in WT (G1 to G10, from low to high). Violin plots display the Log_2_-transformed expression levels within each group, with the orange line indicating the overall trend across transcriptional bins. F. H2A.Z enrichment levels across the ten groups in WT cells. y axis: H2A.Z enrichment level. x axis: 10 quantiles of genes rank by their gene expression levels in WT (G1 to G10, from low to high). Violin plots show the H2A.Z enrichment level within each group, with the orange line indicating the median trend. G. Correlation analysis between transcription activity and H2A.Z enrichment levels. Each dot represents one gene. A strong positive correlation was observed, with the red line indicating the linear regression fit.

To further characterize H2A.Z’s genomic association, we reanalyzed the published genome-wide chromatin profiling of H2A.Z (Wang et al 2025), which demonstrated that H2A.Z was preferentially localized within gene bodies rather than intergenic regions, with pronounced enrichment in coding sequences (CDS) compared to untranslated regions (UTR) (Fig. 1B). Strikingly, ∼76.03% of total H2A.Z peaks were concentrated near the 5’ ends of genes (Fig. 1C-D), suggesting a possible functional link between H2A.Z deposition and transcription.

To test whether H2A.Z distribution correlates with transcriptional activity, we stratified genes into 10 groups based on their expression levels (Fig. 1E). Intriguingly, H2A.Z occupancy progressively increased with elevated gene expression (Fig. 1F), and a genome-wide correlation analysis confirmed a strong positive association (Pearson’s R = 0.44) between H2A.Z enrichment and transcriptional output (Fig. 1G).

Together, these findings demonstrate that H2A.Z is preferentially deposited at transcriptionally active loci in *Tetrahymena*, with a particular bias toward the 5’ regions of highly expressed genes, reinforcing its potential role in transcriptional regulation.

### Bromodomain protein IBD1 associates with SWR Complex

The specific genomic distribution of H2A.Z is fundamental to its regulatory function, and its incorporation into chromatin is mediated by the evolutionarily conserved SWR complex. To investigate the spatial and functional relationship between the SWR complex and H2A.Z in *Tetrahymena*, we characterized the distribution of ARP6 (Table S1), a core SWR subunit. We generated *Tetrahymena* cells expressing endogenously N-terminally HA-tagged ARP6 (ARP6-NHA) (Fig. S1A) and observed by IF that ARP6 exclusively localized to the transcriptionally active MAC (Fig. S1B), mirroring H2A.Z’s nuclear compartmentalization (Fig. 1A).

Genome-wide profiling revealed that ARP6 and H2A.Z exhibited highly correlated genomic distributions (Fig. 2A). ARP6 exhibits strikingly similar enrichment patterns to H2A.Z, with pronounced accumulation in gene bodies rather than intergenic regions, preferential localization to CDS, and a strong 5’ bias (∼64.44% of signal) (Fig. 2B-D). Importantly, >93.5% of ARP6-enriched genes (ARP6+) also displayed H2A.Z occupancy (H2A.Z+) (Fig. 2E). Moreover, we observed mutual reinforcement between the two: H2A.Z+ genes exhibited significantly higher ARP6 enrichment than H2A.Z-genes, and vice versa (Fig. 2F). Consistent with this, their genome-wide enrichment profiles showed a strong positive correlation (Pearson’s R = 0.63) (Fig. 2G), supporting the notion that ARP6, as a component of the SWR complex, mediates H2A.Z deposition in *Tetrahymena*.

**Fig. 2.**
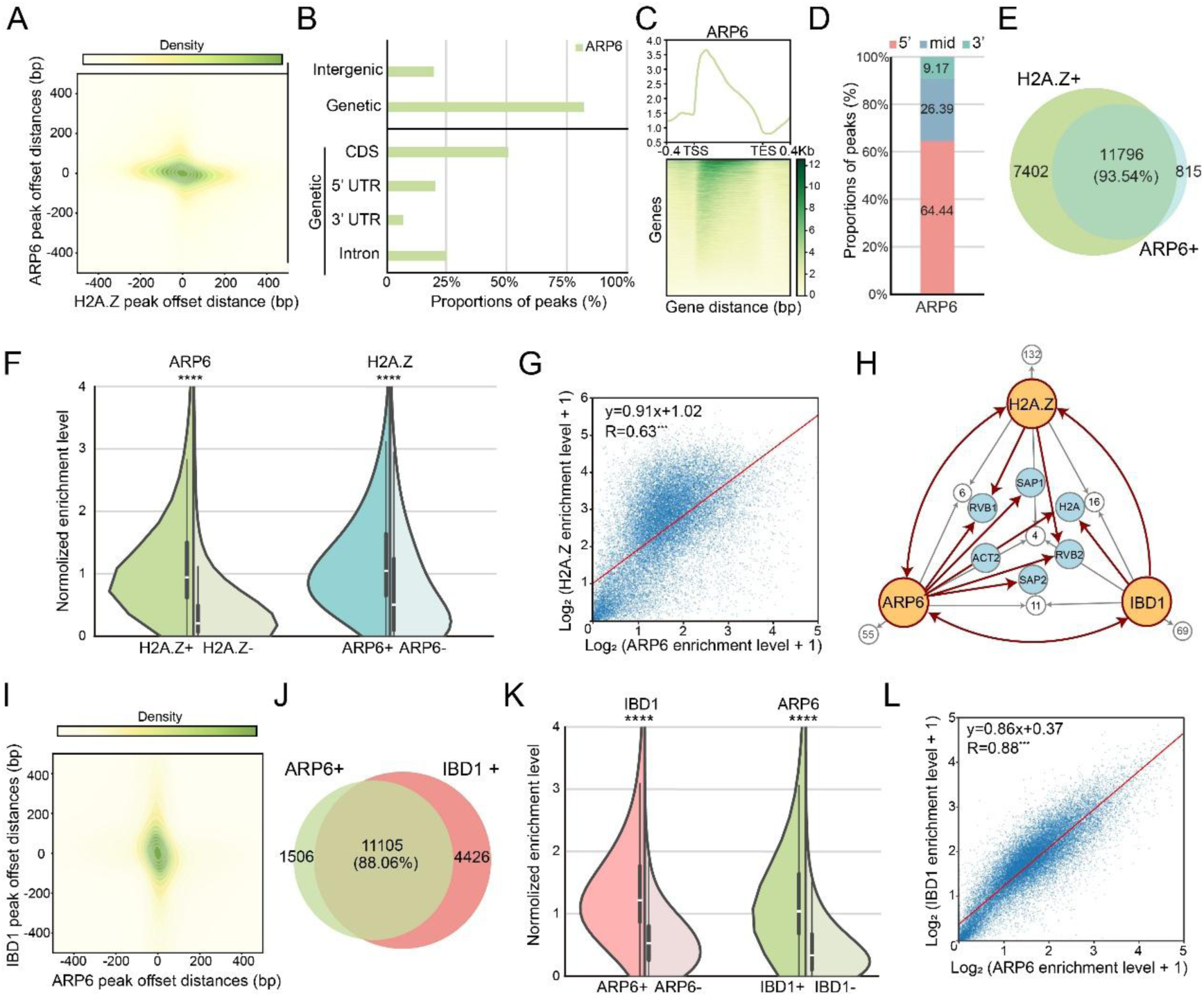
Bromodomain protein IBD1 associates with SWR Complex. A. Analysis of peak offset distances between H2A.Z and ARP6 signals reveals a high degree of overlap, as both x- and y-axis offsets approach zero. The midpoint between adjacent H2A.Z and ARP6 peak summits was determined with the distances from the H2A.Z and ARP6 summits to the midpoint used as the x- and y-axis values, respectively. B. Distribution of ARP6 in the different genomic regions in WT cells. C. Distribution of ARP6 along the gene body shows that it was preferentially enriched at the 5’ end of gene bodies in WT cells. D. Proportions of ARP6 peaks distributed across different gene body regions. E. The overlap between H2A.Z enriched (H2A.Z+) and ARP6 enriched (ARP6+) genes. Genes with high-confident peaks were specifically designated as H2A.Z+ and ARP6+ genes. A significant overlap (93.54%) exists between these two gene sets. F. ARP6 and H2A.Z enrichment levels are significantly enhanced depending on the presence of each other. Violin plots depict the normalized enrichment levels of the other protein on genes upon depletion or enrichment of one protein. G. Correlation analysis between ARP6 and H2A.Z enrichment levels. H. IP-MS interaction network using H2A.Z, ARP6, and IBD1 as baits, respectively. Arrows indicate the direction of bait-prey relationships. Target Proteins are highlighted in blue circles. Gray circles represent the number of additional interactors which not shown individually. I. Analysis of peak offset distances between ARP6 and IBD1 signals reveals a high degree of overlap, as both x- and y-axis offsets approach zero. J. The overlap between ARP6+ and IBD1+ genes. K. IBD1 and ARP6 enrichment levels are significantly enhanced depending on the presence of each other. L. Correlation analysis between ARP6 and IBD1 enrichment levels.

To identify potential regulators of this process, we performed ARP6-NHA immunoprecipitation coupled with mass spectrometry (IP-MS) (Table S2), which revealed interactions with multiple components of the SWR complex— including RVB1, RVB2, ACT2, SAP1, and SAP2—as well as with H2A.Z and canonical H2A (Fig. 2H). Consistent with the role of ARP6 in H2A.Z deposition, ARP6, RVB1, and RVB2 were also identified in the reciprocal H2A.Z-NHA IP-MS (Fig. 2H, Table S2). Notably, ARP6 was found to interact with the bromodomain-containing protein IBD1 (Table S1). Reciprocal IBD1-CHA IP-MS confirmed this association and additionally pulled down H2A.Z and H2A (Fig. S2A, Fig. 2H, Table S2). Genome-wide profiling revealed that IBD1 exhibited pronounced co-localization with ARP6 (Fig. 2I), with approximately 88.06% of IBD1-enriched (IBD1+) genes also showing ARP6-enrichment (ARP6+) (Fig. 2J). We further observed reciprocal reinforcement between IBD1 and ARP6: genes marked by IBD1 displayed significantly higher ARP6 levels than IBD1-genes, and vice versa (Fig. 2K). Moreover, their genome-wide occupancy patterns were strongly correlated (Pearson’s R = 0.88) (Fig. 2L). These findings showed a close spatial association between IBD1 and ARP6, indicating a possible functional link between IBD1 and H2A.Z deposition.

### IBD1 exhibits spatiotemporal co-localization and mutual reinforcement with H2A.Z

To elucidate the functional relationship between IBD1 and H2A.Z, we first examined their expression dynamics and cellular localization. Transcriptome analysis revealed highly synchronous expression of IBD1 and H2A.Z during both vegetative growth and conjugation (C4-C14) (Fig. 3A, Table S1) (Miao et al 2009, Xiong et al 2011). IF staining confirmed that IBD1, like H2A.Z, is exclusively localized to the transcriptionally active MAC and absent from the transcriptionally inert MIC (Fig. 3B, Fig. S2B), supporting a potential functional interplay. Additionally, co-immunoprecipitation followed by western blotting (Co-IP-WB) further confirmed the interaction between IBD1 and H2A.Z (Fig. 3C).

**Fig. 3.**
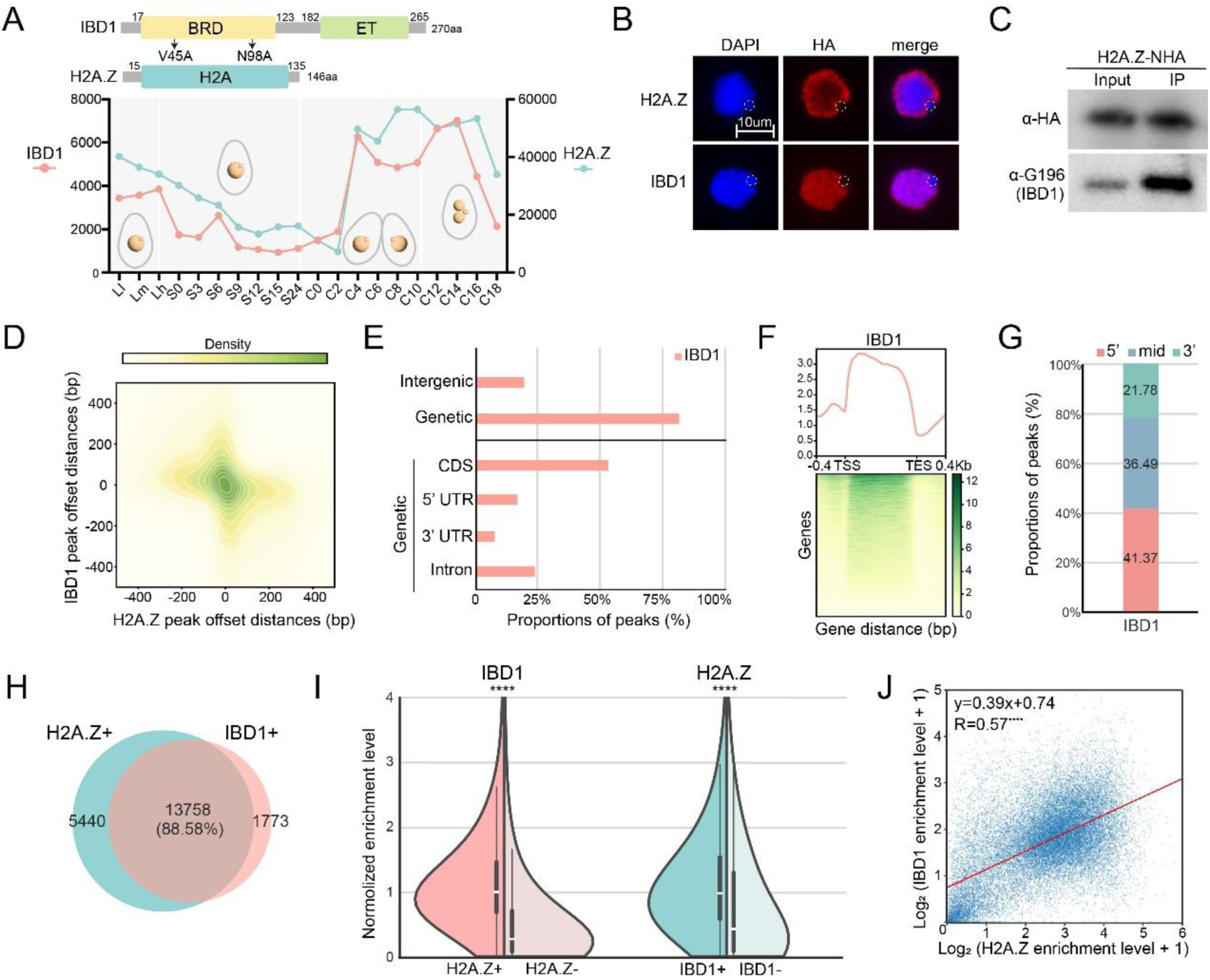
IBD1 exhibits spatiotemporal co-localization and mutual reinforcement with H2A.Z. A. Domain architectures and expression profiles of IBD1 and H2A.Z. The top panel shows the domain structure of IBD1 (with BRD and ET domains) and H2A.Z (with H2A core domain). Mutation sites (V45A and N96A) in the BRD domain of IBD1 used for BRD-mutated constructs are indicated. The bottom panel displays the expression profiles of IBD1 (red line) and H2A.Z (blue line) across different developmental stages. B. Cellular localization of HA-tagged H2A.Z and IBD1. Both proteins are exclusively localized to the MAC, with no detectable HA signal in the MIC, which is outlined with dotted circles. Nuclei were counterstained with DAPI (blue). C. IP-WB analysis showing the interaction between H2A.Z and IBD1. H2A.Z-NHA serves as the bait, and IBD1-3 × G196 is detected as the prey using an α-G196 antibody. D. Analysis of peak offset distances between H2A.Z and IBD1 signals reveals a high degree of overlap, as both x-and y-axis offsets approach zero. E. Distribution of IBD1 in the different genomic regions in WT cells. F. Distribution of IBD1 along the gene body shows that it was preferentially enriched at the 5’ end of gene bodies in WT cells. G. Proportions of IBD1 peaks distributed across different gene body regions. H. The overlap between H2A.Z+ and IBD1+ genes. I. IBD1 and H2A.Z enrichment levels are significantly enhanced depending on the presence of each other. J. Correlation analysis between IBD1 and H2A.Z enrichment levels.

Genome-wide profiling demonstrated that IBD1 and H2A.Z exhibit highly correlated genomic distributions (Fig. 3D). Both factors predominantly occupied genic regions rather than intergenic sequences and were particularly enriched at CDS relative to UTRs, with pronounced 5’ bias (Fig. 3E-F). Although H2A.Z exhibited stronger 5’ enrichment (76.03%) than IBD1 (41.37%) (Fig. 1D, Fig. 3G), a substantial proportion of IBD1-bound (IBD1+) genes (88.58%) also showed H2A.Z occupancy (H2A.Z+) (Fig. 3H). Importantly, we observed reciprocal reinforcement between these two factors: genes marked by IBD1 (IBD1+) displayed significantly higher H2A.Z levels than IBD1-genes, and in turn, H2A.Z+ genes showed markedly higher IBD1 occupancy than H2A.Z-genes (Fig. 3I). Furthermore, their genome-wide occupancy patterns were significantly correlated (Pearson’s R = 0.57) (Fig. 3J). These comprehensive analyses established that IBD1 associates with H2A.Z, suggesting its potential role in H2A.Z deposition.

### IBD1 bromodomain guides H2A.Z deposition

To investigate whether IBD1 influences H2A.Z chromatin targeting, we generated Δ*IBD1* cells (Fig. S3A) and analyzed the genomic distributions of ARP6 and H2A.Z in this background. Strikingly, while ARP6 cellular localization and total protein levels remained unchanged (Fig. S3B-C), its 5′-enrichment was significantly attenuated in Δ*IBD1* cells (Fig. 4A). Concomitantly, H2A.Z deposition at these regions was markedly reduced, though its global levels and nuclear distribution were unaffected (Fig. 4B, Fig. S3D-E). These results establish that IBD1 directs ARP6, a core SWR complex subunit, to promote H2A.Z incorporation at gene 5′ ends, revealing a critical IBD1-ARP6-H2A.Z regulatory axis.

**Fig. 4.**
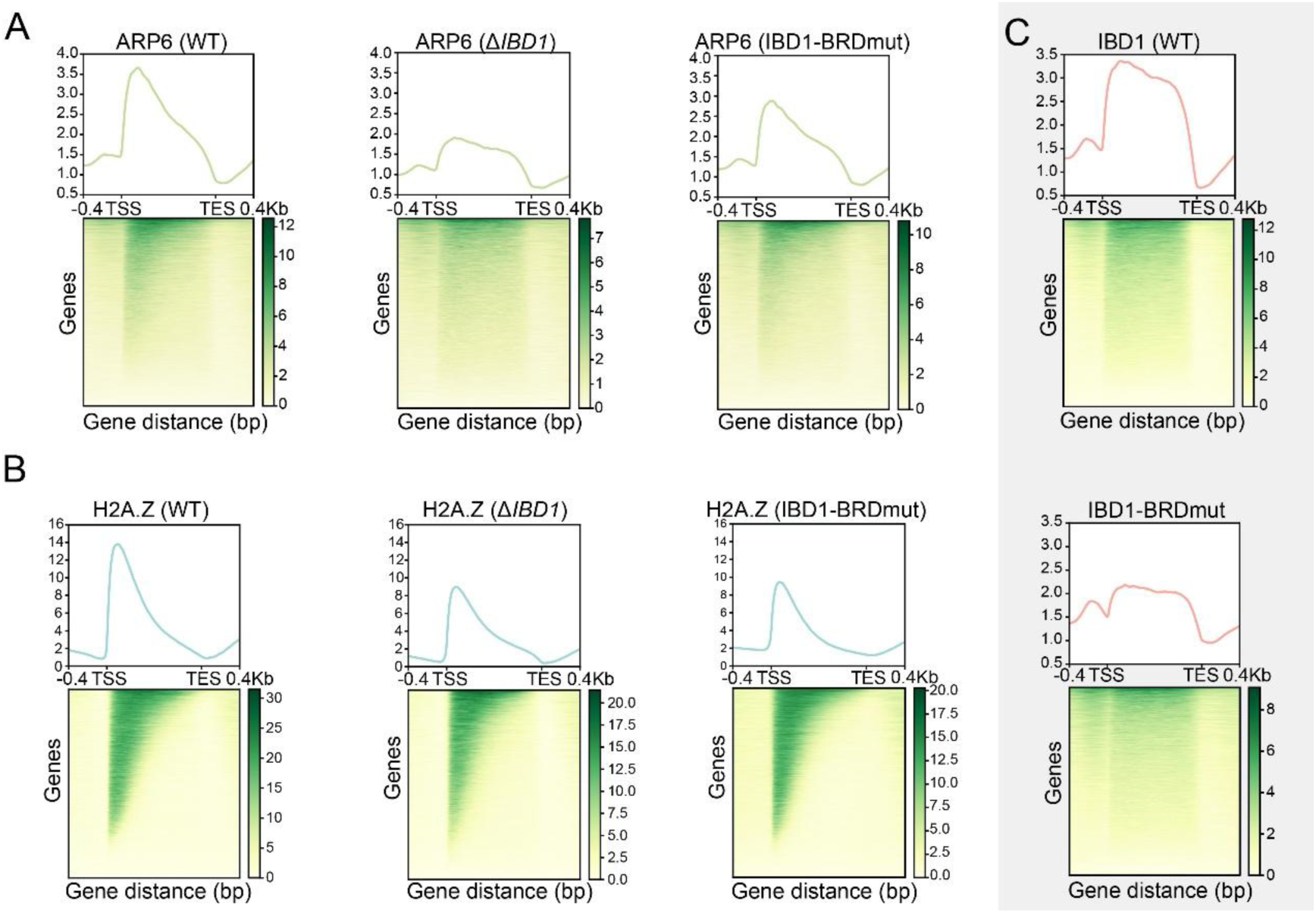
IBD1 bromodomain guides H2A.Z deposition. A. The enrichment of ARP6 at the 5′ ends of gene bodies is reduced in both Δ*IBD1* and IBD1-BRDmut cells, compared with that in WT. B. The enrichment of H2A.Z at the 5′ ends of gene bodies is reduced in both Δ*IBD1* and IBD1-BRDmut cells, compared with that in WT. C. The enrichment of IBD1 at the 5′ ends of gene bodies is reduced in both Δ*IBD1* and IBD1-BRDmut cells, compared with that in WT.

IBD1, a conserved BET family member, harbors an N-terminal bromodomain (BRD) and a C-terminal extraterminal (ET) domain (Fig. 3A). Sequence alignment of the IBD1 BRD with human BET family orthologs demonstrated conservation of key acetyl-lysine (Kac)-binding residues (Fig. S3F), suggesting functional recognition of histone acetylation marks. To test whether BRD-mediated histone engagement governs H2A.Z deposition, we generated IBD1-BRD mutants (V45A/N98A; IBD1-BRDmut) impairing Kac binding (Fig. S3G). Genome-wide profiling revealed significantly reduced 5′ occupancy of IBD1 in IBD1-BRDmut cells (Fig. 4C), despite unaffected cellular localization and increased total protein level (Fig. S3H-I). Crucially, both ARP6 and H2A.Z reduced their 5′ enrichment in IBD1-BRDmut cells, phenocopying Δ*IBD1* phenotypes (Fig. 4A-B). ARP6 and H2A.Z localization and protein levels remained unaltered (Fig. S3B-E), confirming that the observed defect results from impaired BRD-dependent chromatin recruitment rather than altered protein turnover. Collectively, these data demonstrate that the BRD of IBD1 is essential for ARP6 recruitment and the subsequent H2A.Z deposition at gene 5′ ends, directly linking histone acetylation recognition to H2A.Z deposition.

### Histone acetylation governs H2A.Z chromatin deposition through BRD-mediated recognition

Having established the essential role of IBD1’s BRD in chromatin association, we next aimed to delineate its specific histone acetylation recognition profile and mechanistic role in H2A.Z deposition. Previous characterization through histone peptide arrays revealed that recombinant *Tetrahymena* IBD1 preferentially binds H3 peptides di-acetylated at lysines 9 and 14 (H3K9/14ac) (Saettone et al 2018). We performed quantitative binding assays with purified wild-type IBD1 protein and confirmed its strongest affinity for H3K9/14ac peptides, with significantly weaker binding to mono-acetylated or unmodified controls (Fig. 5A, Table S3). This interaction specificity was completely abolished in IBD1-BRDmut protein (Fig. 5A), demonstrating the BRD’s exclusive responsibility for H3K9/14ac recognition.

**Fig. 5.**
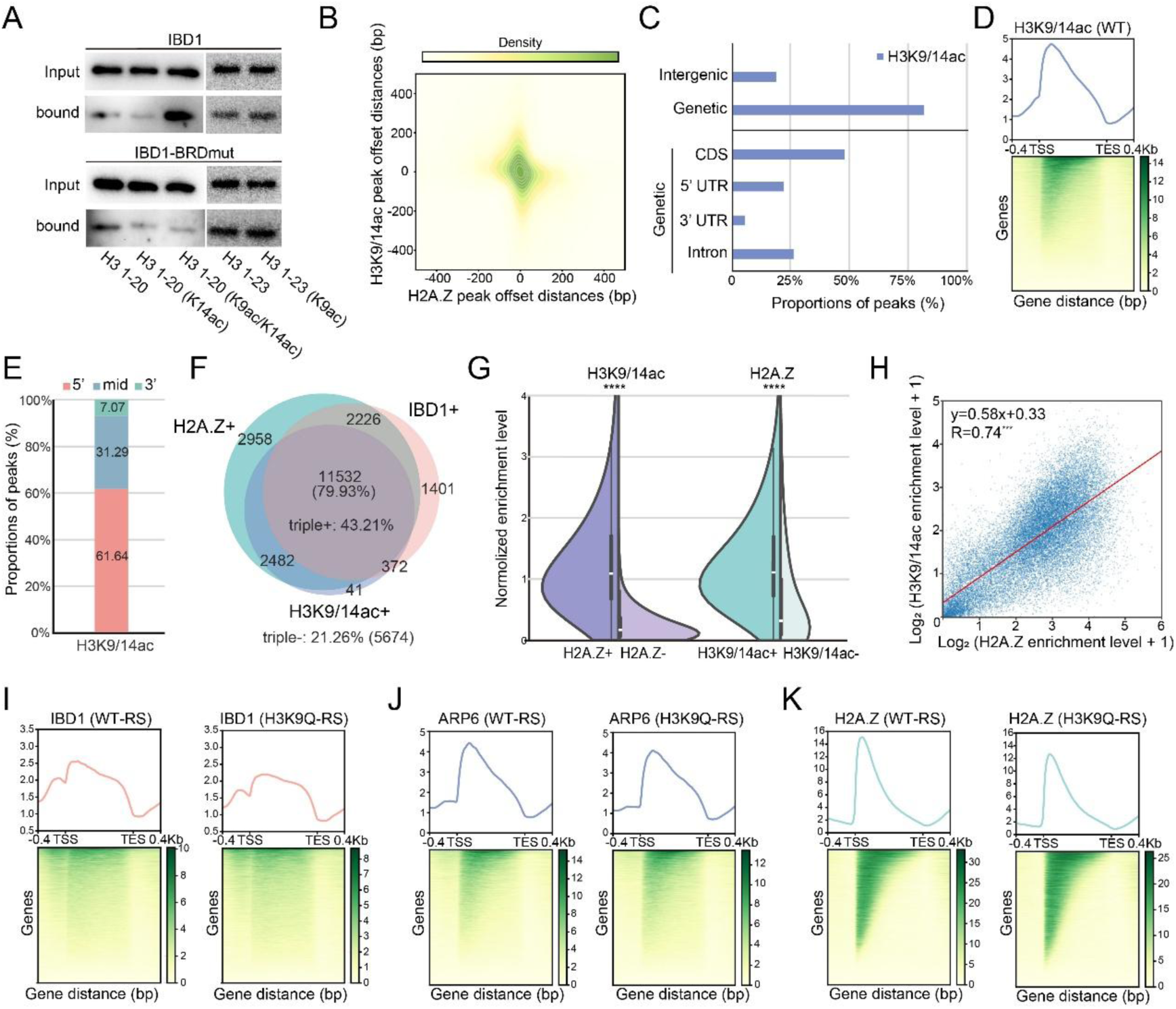
Histone acetylation governs H2A.Z chromatin deposition through BRD-mediated recognition. A. Peptide binding assay showing the interaction of purified IBD1 and IBD1-BRDmut proteins with histone H3 tail peptides carrying different modifications. B. Analysis of peak offset distances between H2A.Z and H3K9/14ac signals reveals a high degree of overlap, as both x- and y-axis offsets approach zero. C. Distribution of H3K9/14ac in the different genomic regions in WT cells. D. Distribution of H2A.Z along the gene body shows that it was preferentially enriched at the 5’ end of gene bodies in WT cells. E. Proportions of IBD1 peaks distributed across different gene body regions. F. The overlap between H2A.Z+, IBD1+ and H3K9/14ac+ genes. G. H3K9/14ac and H2A.Z enrichment levels are significantly enhanced depending on the presence of each other. H. Correlation analysis between H2A.Z and H3K9/14ac enrichment levels. I. The enrichment of IBD1 at the 5′ ends of gene bodies is reduced in H3K9Q-RS cells, compared with that in WT. J. The enrichment of ARP6 at the 5′ ends of gene bodies is reduced in H3K9Q-RS cells, compared with that in WT. K. The enrichment of H2A.Z at the 5′ ends of gene bodies is reduced in H3K9Q-RS cells, compared with that in WT.

To assess the *in vivo* relevance of IBD1-associated chromatin regulation, we performed IF staining and observed that H3K9/14ac is exclusively localized to the MAC, consistent with the nuclear distribution of IBD1 (Fig. S4A). Genome-wide profiling revealed that H3K9/14ac, IBD1, and H2A.Z share highly correlated genomic distributions, with all three factors exhibiting a strong preference for genic regions, particularly coding sequences (CDSs) (Fig. S4B, Fig. 5B-C). Notably, H3K9/14ac exhibited pronounced co-localization with IBD1 and H2A.Z at gene 5′ ends (Fig. 5D-E), with ∼79.93% of H3K9/14ac-enriched genes also showing co-enrichment of IBD1 and H2A.Z (Fig. 5F). We further observed reciprocal reinforcement among all three factors: genes marked by IBD1 (IBD1+) showed significantly higher H3K9/14ac levels than IBD1-genes, and vice versa (Fig. S4C). Similarly, H3K9/14ac+ genes exhibited greater H2A.Z deposition than H3K9/14ac-genes, and vice versa (Fig. 5G). Quantification revealed strong positive correlation between H3K9/14ac levels and both IBD1 occupancy (Pearson’s R = 0.77) (Fig. S4D) and H2A.Z deposition (R=0.74) (Fig. 5H). We also measured H3K9/K14ac levels following the deletion or mutation of *IBD1* gene, but not many changes were observed, including cellular localization, protein level and genomic distribution (Fig. S4E-G).

To validate the effect of H3K9/K14ac in H2A.Z deposition, we next generated an H3K9Q mutation that mimics constitutive deacetylation at this residue to directly test the role of histone acetylation *in vivo* (Fig. S4H). While attempts to create an H3K14Q mutant were unsuccessful due to viability constraints in *Tetrahymena*, the H3K9Q mutation alone caused a moderate reduction in 5’ enrichment of all three marks (IBD1, ARP6 and H2A.Z) compared to wild-type controls (Fig. 5I-K), without affecting their subcellular localization or protein levels (Fig. S4I-N). These results confirm that H3K9 acetylation affects proper H2A.Z occupancy *in vivo*.

Together, our findings establish a molecular pathway wherein IBD1 is recruited to chromatin through BRD-dependent recognition of histone acetylation, specifically H3K9/14ac marks, subsequently guiding the SWR complex to mediate H2A.Z deposition at gene 5’ ends. This mechanism ensures the precise genomic occupancy of H2A.Z.

### H2A.Z cooperates with histone acetylation to fine-tune transcription and prevent hyperactivation

The interplay between histone acetylation and H2A.Z deposition may constitute a pivotal epigenetic regulatory mechanism in transcription. To elucidate the functional relationship within the histone acetylation-IBD1-H2A.Z axis, we first analyzed transcriptomes of Δ*IBD1* and IBD1-BRDmut cells. Strikingly, both mutants triggered widespread transcriptional upregulation (Fig. 6A-B), with highly overlapping sets of upregulated and downregulated genes (Fig. S5A-B). This transcriptional activation was accompanied by a global reduction in H2A.Z occupancy (Fig. 4B), suggesting a repressive role for H2A.Z in transcriptional control.

**Fig. 6.**
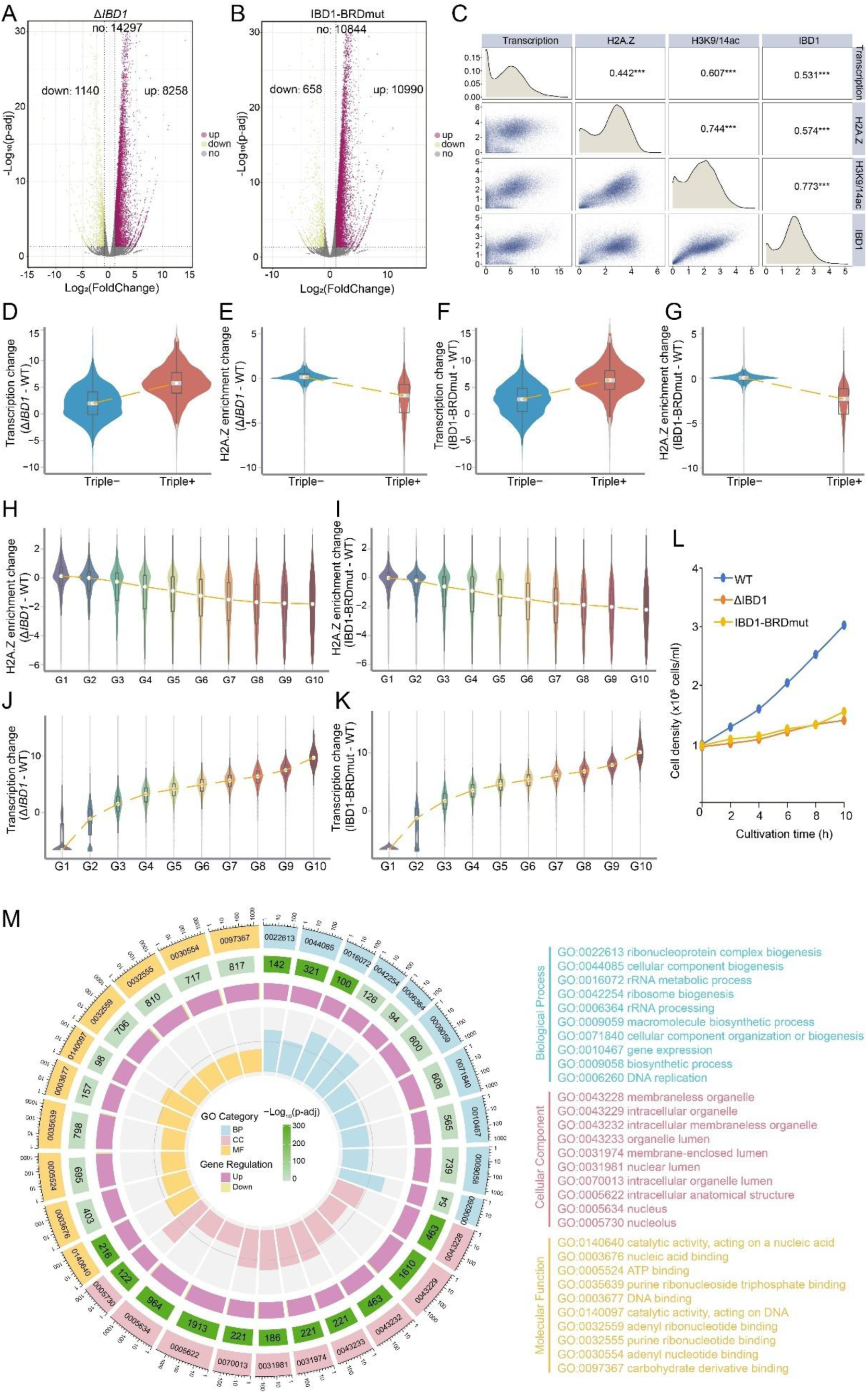
H2A.Z cooperates with histone acetylation to fine-tune transcription and prevent hyperactivation. A. Volcano plot showing differential gene expression in the Δ*IBD1* cells compared to that in WT. The x-axis represents the log_2_(Fold Change), and the y-axis shows the –log_10_(p-adj). Genes with significantly increased expression are shown in magenta; genes with significantly decreased expression are shown in yellow; and non-significant genes are shown in gray. B. Volcano plot showing differential gene expression in the IBD1-BRDmut cells compared to WT. C. Correlations between H2A.Z, IBD1, H3K9/K14ac, and gene expression levels. H2A.Z, IBD1, H3K9/K14ac levels: the average enrichment score over gene bodies, corrected using background genes, calculated with computeMatrix and custom scripts. Gene expression levels: FPKM for RNA-seq of WT. D. Changes in transcriptional activity of genes co-marked by H2A.Z, IBD1, and H3K9/14ac (triple+) and genes lacking all three marks (triple-) in Δ*IBD1* cells. Violin plots show the distribution of transcriptional changes between triple+ and triple-genes. E. Changes in H2A.Z enrichment of triple+ and triple-genes in the Δ*IBD1* cells. F. Changes in transcriptional activity of triple+ and triple-genes in the IBD1-BRDmut cells. G. Changes in H2A.Z enrichment of triple+ and triple-genes in the IBD1-BRDmut cells. H. Changes in transcriptional activity across ten gene groups in the Δ*IBD1* cells. y axis: transcription changes (Δ*IBD1-WT*). x axis: 10 quantiles of genes rank by their transcription levels in WT (G1 to G10, from low to high). Violin plots show the normalized transcriptional changes in each group, with the orange line indicating the overall trend. I. Changes in H2A.Z enrichment across ten gene groups in the Δ*IBD1* cells. y axis: H2A.Z enrichment changes (Δ*IBD1-WT*). x axis: 10 quantiles of genes rank by their transcription levels in WT (G1 to G10, from low to high). Violin plots show the H2A.Z normalized enrichment level changes within each group, with the orange line indicating the overall trend. J. Changes in transcriptional activity across ten gene groups in the IBD1-BRDmut cells. K. Changes in H2A.Z enrichment across ten gene groups in the IBD1-BRDmut cells. L. Growth curves of WT, Δ*IBD1*, and IBD1-BRDmut cells. Cell density was measured at the indicated time points during cultivation. M. GO enrichment analysis of shared differentially expressed genes between Δ*IBD1* and IBD1-BRDmut strains. The circular plot illustrates significantly enriched Gene Ontology (GO) terms, categorized into Biological Process (BP, blue), Cellular Component (CC, pink), and Molecular Function (MF, yellow). For each category, the top 10 GO terms with the lowest adjusted p-values are shown. Each sector represents a GO term, with arc length proportional to the log_10_-transformed number of background genes. From inside to outside: (1) GO category background with term labels and coordinate axis (log_10_ scale); (2) segment length represents enriched gene counts (log10-transformed), colored by −log_10_(adjusted p-value); (3) proportion of upregulated (purple) and downregulated (yellow) genes; (4) segment height represents the Rich Factor (enriched genes/background genes), with color matching GO categories.

To delineate the regulatory logic, we performed genome-wide correlation analyses and observed strong positive associations among H2A.Z, IBD1, H3K9/K14ac, and Pol II-transcribed gene expression (Fig. 6C). To explore the relationship between H2A.Z and histone acetylation, we focused on genes marked by H3K9/14ac. Among these H3K9/14ac+ genes, those that were also marked by H2A.Z exhibited significantly higher H3K9/14ac enrichment levels compared to H2A.Z-genes (Fig. S5C). Moreover, H2A.Z and H3K9/14ac enrichment levels are positively correlated, and both are associated with higher transcriptional activity (Fig. S5D). Notably, most WT genes were either co-enriched (triple+: 43.21%) or co-depleted (triple-: 21.26%) for all three marks (H2A.Z, IBD1, and H3K9/K14ac) (Fig. 5F). Critically, in both Δ*IBD1* and IBD1-BRDmut cells, H2A.Z levels at triple+ genes were substantially diminished (Fig. 6D-E), concomitant with elevated transcription (Fig. 6F-G), whereas triple-genes remained unaffected (Fig. 6D-G). Given comparable H3K9/K14ac levels (Fig. S5E-F), we conclude that transcriptional upregulation is driven primarily by H2A.Z loss.

To further dissect this relationship, we stratified WT-expressed genes into 10 quantiles by expression level. While H3K9/K14ac levels were unchanged across all groups in mutants (Fig. S5G-H), H2A.Z depletion was most pronounced in highly expressed genes (Fig. 6H-I), which also exhibited the greatest transcriptional elevation (Fig. 6J-K). Notably, H2A.Z reduction correlated with transcriptional increase in Δ*IBD1*/IBD1-BRDmut cells (Pearson’s R = −0.267, −0.296) (Fig. S5I-J), supporting H2A.Z’s role as a transcriptional dampener at highly active loci.

This transcriptional dysregulation led to profound phenotypic consequences: both mutants exhibited growth defects (Fig. 6L) and widespread transcriptional hyperactivation in Gene Ontology (GO) categories, with the most pronounced effects observed in the cellular component category (Fig. 6M). These defects aligned with global H2A.Z loss and aberrant gene activation, demonstrating that the histone acetylation-IBD1-H2A.Z axis is essential for maintaining transcriptional homeostasis and cellular fitness.

## Discussion

In this study, we uncover a critical regulatory axis in which the bromodomain protein IBD1 recognizes histone acetylation, specifically H3K9/14 di-acetylation to direct SWR complex-mediated H2A.Z deposition, ensuring precise transcriptional control in *Tetrahymena* (Fig. 7). Our findings reveal that disruption of this pathway, either through loss of IBD1 or impairment of its bromodomain, leads to aberrant H2A.Z occupancy and transcriptional hyperactivation, highlighting the dual role of H2A.Z in both facilitating basal transcription and preventing excessive gene expression.

**Fig. 7.**
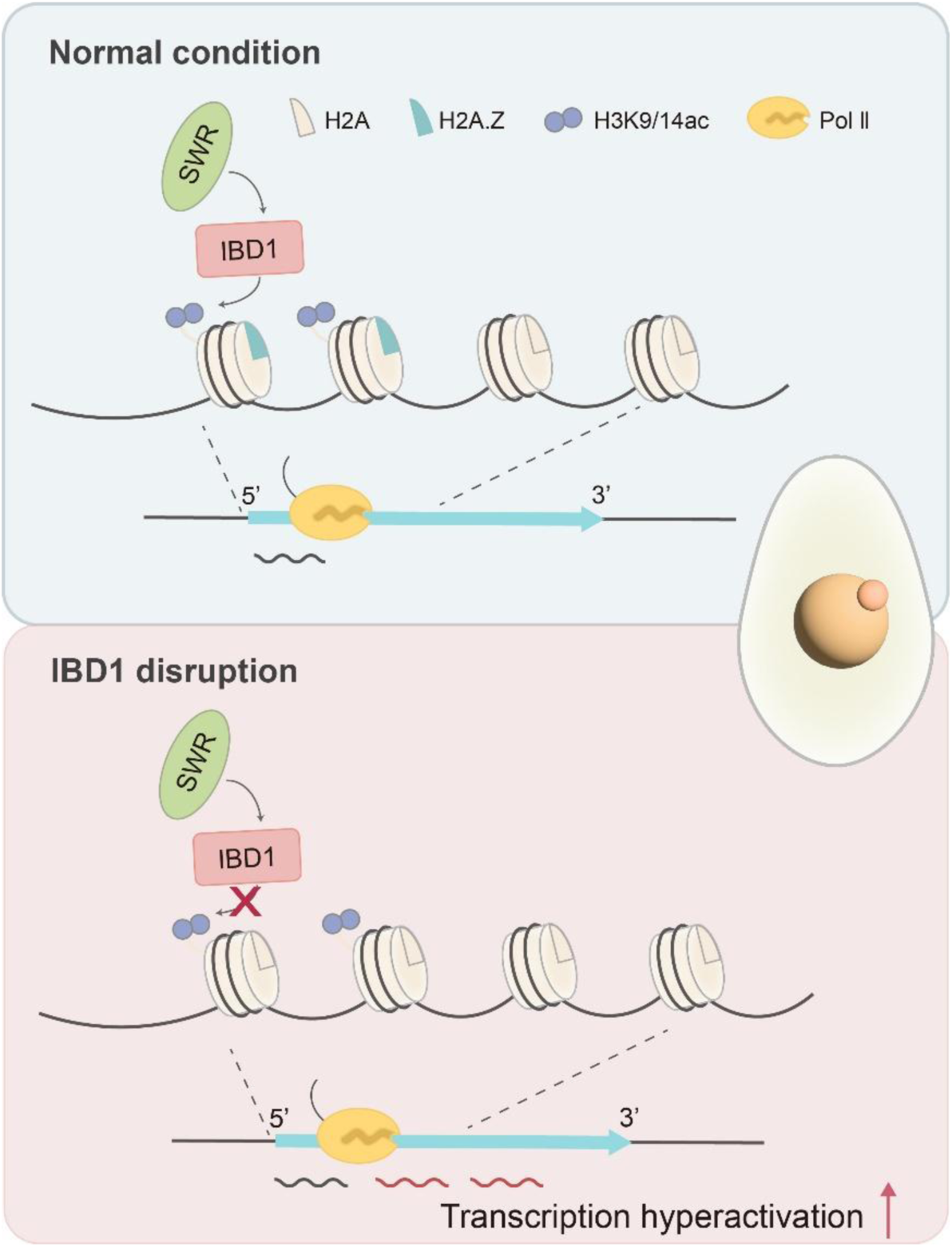
IBD1 bridges histone acetylation and H2A.Z deposition to regulate transcription. Top panel (Normal condition): IBD1 recognizes H3K9/14 acetylation and recruits the SWR complex to deposit H2A.Z at the 5′ ends of active genes, thereby maintaining proper transcriptional activity. Bottom panel (IBD1 disruption): Disruption of IBD1 impairs H2A.Z deposition despite the presence of histone acetylation, resulting in transcriptional hyperactivation.

### IBD1 bridges histone acetylation and H2A.Z deposition

Our work identifies IBD1 as a key adaptor that selectively binds H3K9/14ac and recruits the SWR complex, ensuring localized H2A.Z deposition. This finding aligns with recent reports in mammals showing that the YEATS domain of GAS41 recognizes H3K14ac/H3K27ac to promote H2A.Z occupancy (Kikuchi et al 2023), suggesting an evolutionarily conserved mechanism linking histone acetylation to H2A.Z dynamics. However, unlike GAS41, which primarily functions in transcriptional activation, IBD1 appears to play a role in restraining transcription, as its loss leads to hyperactivation. This distinction underscores the context-dependent nature of H2A.Z regulation, where its functional outcome may depend on associated histone marks and the specific reader proteins involved.

Notably, while IBD1 exhibits a strong preference for H3K9/K14 double acetylation, it also shows affinity for H2Aac and H4ac (Saettone et al 2018), suggesting that other histone acetylation marks may contribute to IBD1-mediated H2A.Z distribution. This raises the possibility that IBD1 integrates multiple acetylation signals to fine-tune H2A.Z incorporation, a hypothesis that warrants further investigation. Additionally, *Tetrahymena* encodes 14 BRD-containing proteins, some of which may also participate in H2A.Z regulation (Saettone et al 2018, Zhang et al 2025). Future studies should explore the potential roles of these BRD proteins in chromatin remodeling and H2A.Z dynamics, as well as their functional redundancy or specialization.

### H2A.Z as a transcriptional buffer and epigenetic integrator

Our discovery that disruption of the IBD1-H2A.Z axis leads to transcriptional hyperactivation challenges the conventional view of H2A.Z as a purely activating mark. While H2A.Z is indeed associated with transcriptionally active features, including H3K4me3 and 6mA in *Tetrahymena*, which similarly accumulate at 5’ ends of Pol II-transcribed genes and are restricted to the transcriptionally active MAC (Wang et al 2025). Our findings reveal a more sophisticated regulatory role. Rather than simply promoting transcription, H2A.Z appears to function as a transcriptional buffer, maintaining optimal expression levels while preventing aberrant overexpression. This repressive function in highly acetylated regions is particularly significant, suggesting H2A.Z acts as a molecular “governor” that constrains transcriptional output despite strong activating signals. The dual regulatory capacity of H2A.Z mirrors observations in yeast and mammalian systems, where it participates in both activation and silencing (Diegmüller et al 2025, Wong & Tremethick 2025), highlighting an evolutionarily conserved mechanism whose outcome depends on chromatin context.

This functional versatility likely stems from H2A.Z’s complex interplay with other PTMs. Hyperacetylation of its N-terminal lysines correlates with activation, while monoubiquitination of C-terminal residues links to repression, and SUMOylation has been associated with silencing (Bruce et al 2005, Fukuto et al 2018, Surface et al 2016). Within this framework, the histone acetylation-IBD1-H2A.Z axis represents a sophisticated regulatory circuit that fine-tunes gene expression, establishing negative feedback at highly active loci to prevent transcriptional runaway. The complexity of this regulation suggests additional layers of control remain to be discovered, particularly regarding how distinct H2A.Z modification patterns are established, interpreted by reader proteins, and integrated with other epigenetic signals. Future studies systematically mapping H2A.Z modification states across transcriptional conditions and characterizing their crosstalk with other chromatin features will not only clarify H2A.Z’s dual roles but also advance our understanding of how eukaryotic cells achieve precise transcriptional control through dynamic chromatin regulation.

### Broader implications for epigenetic regulation

Our findings significantly expand the known functional repertoire of BRD proteins in chromatin biology. While BRD proteins like IBD1 are well-established as “readers” of acetylated histones, their direct involvement in guiding H2A.Z deposition through coordination with the SWR complex reveals a previously underappreciated layer of epigenetic regulation. This discovery not only bridges the gap between histone acetylation signals and chromatin remodeling but also suggests that BRD proteins may serve as critical adaptors that integrate multiple epigenetic pathways. The evolutionary conservation of these mechanisms across eukaryotes implies they represent a fundamental aspect of chromatin-based regulation, with potentially wide-ranging implications for understanding how cells interpret and respond to epigenetic cues in various biological contexts.

The dynamic interplay between histone acetylation and H2A.Z deposition we observed points to the existence of sophisticated feedback mechanisms in chromatin regulation. The incorporation of H2A.Z, once established through acetylation-dependent recruitment, may itself influence subsequent transcriptional output, maintaining chromatin state homeostasis. This loop becomes particularly intriguing when considering how it might interface with other histone modifications, such as methylation, to fine-tune IBD1 targeting and H2A.Z dynamics. Such multi-layered regulation likely plays crucial roles in processes requiring rapid transcriptional adjustments, including cellular stress responses and developmental transitions, where precise control of gene expression patterns is essential. Future investigations into how this regulatory axis operates in different genomic and cellular contexts will not only enhance our understanding of epigenetic memory mechanisms but may also reveal new principles governing transcriptional plasticity in both normal physiology and disease states.

## Materials and Methods

### Cell culture

Wild-type SB210 and CU428 cells of *Tetrahymena thermophila* were obtained from the *Tetrahymena* Stock Center (http://tetrahymena.vet.cornell.edu). Cells were cultured in SPP medium at 30°C, 200 rpm, to the logarithmic phase (2-3 × 10^5^ cells/mL) for subsequent experiments, following standard protocols (Cassidy-Hanley 2012). For starvation, cells were transferred to 10 mM Tris-HCl (pH 7.4) and incubated overnight at 30°C. Cell density was measured using a Beckman Coulter Z2 Particle Counter.

### Generation of *Tetrahymena* cells

The construction of the cells was conducted according to established protocols (Chalker 2012). For the Δ*IBD1* cells, the IBD1 coding sequence (CDS) was replaced by a *neo4* cassette (Mochizuki et al 2008) (Fig. S3A). Paromomycin (Sigma, P5057) concentration was increased to select for the fully knockout cells. For the epitope-tagging cells, the protein was tagged with either an HA or a 3 × G196 tag at the N-terminus or C-terminus (Tatsumi et al 2017). Additionally, the *chx* cassette, *bsr2* cassette or *neo4* cassette was inserted into the 5’ or 3’ flanking regions of the gene (Mochizuki et al 2008, Qiao et al 2022, Wang et al 2019) (Fig. S1A, Fig. S2A, S3G). The concentrations of cycloheximide (for *chx*-containing constructs), paromomycin (for *neo4*-containing constructs) or blasticidin S (Solarbio, B9300) (for *bsr2*-containing constructs) were increased to select for the complete replacement cells. The construction of the H3H4-RS and H3K9QH4-RS cells was carried out based on a previous study (Liu et al 2004) (Fig. S4H). All the detailed information about the cells and primers used for cell construction is available in the supplementary information (Table S4-S5).

### Immunofluorescence staining

Immunofluorescence (IF) staining was performed according to a previously study (Gao et al 2013). Cells were fixed using 2% paraformaldehyde (Sigma, P6148) and permeabilized with 0.4% Triton X-100 (Sigma, T8787). The primary antibody was α-HA (Cell Signaling, 3724) or α-H3K9/14ac (Abcam, ab232952), and the secondary antibody was Goat anti-Rabbit IgG (H+L) (Invitrogen, A21428).

### Western blot

Western blot samples were prepared according to previously described protocols (Tian & Loidl 2018). A total of 5 × 10⁵ cells were resuspended in 500 µL of 10% trichloroacetic acid (TCA) and incubated on ice for 30 minutes. After centrifugation, the pellet was resuspended in SDS loading buffer for subsequent Western blot analysis. Primary antibodies included α-HA (Cell Signaling, C29F4), α-G196 (Funakoshi, R-G-001), α-H3K9/14ac (Abcam, ab232952), and α-tubulin (Millipore, CP06). The secondary antibody was Goat anti-Rabbit IgG (H+L) HRP Conjugated (TransGen Biotech, HS101) or Goat anti-Mouse IgG (H+L) HRP Conjugated (TransGen Biotech, HS201).

### Immunoprecipitation (IP) and mass spectrometry analysis

Immunoprecipitation (IP) was performed following the procedure outlined in a previous study (Wang et al 2025). After the elution using HA peptide (Sigma, I2149), the protein pellet was analyzed by mass spectrometry at the Analysis Center of Agrobiology and Environmental Sciences, Zhejiang University or dissolution in 1 × SDS loading buffer for Western blot. IP-MS results were analyzed using SAINTexpress v3.6.3 (Teo et al 2014), with peptide-spectrum matches (PSMs) as input and wild-type cells as the negative control. Proteins with a false discovery rate (FDR) ≤ 0.05 were defined as high-confidence interactors. The original IP-MS datasets are available in Supplementary Data 1.

### Recombinant protein purification

Codon-optimized full-length IBD1 and its bromodomain mutant were cloned into the pET28a (+) vector with an N-terminal 6 × His tag (Tsingke Biotechnology Co., Ltd.). The plasmids were transformed into *E. coli* BL21 (DE3) (TransGen Biotech, CD701), cultured at 37°C to the optimal density, and induced with 0.25 mM IPTG at 16°C for 18 hours. Cells were harvested, resuspended in lysis buffer (50 mM Tris-HCl, pH 7.8, 300 mM NaCl, 20 mM imidazole, 10% glycerol), and disrupted using a high-pressure homogenizer. After centrifugation, the supernatant was incubated with Ni Sepharose (Cytiva, 17531801) for affinity purification. Bound proteins were eluted with buffer (20 mM Tris-HCl, pH 8.0, 300 mM NaCl, 200 mM imidazole, 2 mM DTT) and further purified on a Heparin HP column (Cytiva, 17040701), eluting with buffer (20 mM Tris-HCl, pH 7.5, 200 mM NaCl, 2 mM DTT, 5% glycerol).

### Peptide binding assay

A total of 2 µg of protein and 2 µg of peptide were mixed in 1 mL of binding buffer (50 mM Tris-HCl, pH 7.5, 150 mM NaCl, 0.1% NP-40, 50 µM ZnAc, 2mM DTT, and 100 µg/mL BSA), followed by the addition of 30 µL of prewashed streptavidin magnetic beads (Thermo Scientific, 88816). The reaction was incubated with gentle rotation at 4°C for 2 hours. The beads were then washed five times with 1 mL of binding buffer, each wash lasting 5 minutes. Finally, the beads were resuspended in 50 µL 1 × SDS loading buffer and boiled for Western blot analysis. Peptide sequences used in binding assays are listed in the supplementary information (Table S3).

### Genome-wide profiling sample preparation

Genome-wide profiling of H2A.Z was performed using native ChIP, while CUT&Tag was used to profile IBD1, ARP6, and H3K9/14ac.

ChIP-seq samples were prepared following an established protocol (Chen et al 2016, Cuddapah et al 2009, Wang et al 2025). A total of 1 × 10⁸ MACs from HA-tagged cells were digested with Micrococcal Nuclease (NEB, M0247S) to generate mononucleosomes. The supernatant was incubated with anti-HA magnetic beads (Thermo Scientific, 88837) at 4°C for 22 hours to capture HA-tagged proteins bound to DNA. Elution was performed using HA peptide (Sigma, I2149).

Nuclei for CUT&Tag were prepared as follows. A total of 1 × 10⁶ *Tetrahymena* cells were washed once with 10 mM Tris-HCl, pH 7.4, and resuspended in 1 ml lysis buffer containing 10 mM HEPES, pH 7.4, 10 mM KCl, 0.5 mM spermidine, 0.1% Triton X-100, 20% glycerol, 1 × protease inhibitor (PI), and 5 mM sodium butyrate. The suspension was vortexed vigorously for 1 min, followed by incubation on ice for 10 min. The lysate was then centrifuged at 1000 rpm for 5 min, after which the nuclei appeared frosted. The nuclei were subsequently washed once with wash buffer 1 (10 mM HEPES, pH 7.4, 0.5 mM spermidine, 2 mM EDTA, pH 8.0, 150 mM NaCl, 0.1% BSA, 1 × PI, and 5 mM sodium butyrate), followed by an additional wash with wash buffer 2 (10 mM HEPES, pH 7.4, 0.5 mM spermidine, 150 mM NaCl, 0.1% BSA, 1× PI, and 5 mM sodium butyrate). The prepared nuclei were then used for CUT&Tag, following the protocol provided in the kit handbook (Yeasen, 12597ES). The primary antibody used was α-HA (Cell Signaling, 3724) or α-H3K9/14ac (Abcam, ab232952).

### Genome-wide profiling data analysis

All Illumina short-read DNA sequencing data were processed using Trim Galore v0.6.10 for quality control, including adapter removal, filtering of short reads (<36 bp), and trimming of low-quality bases from both ends (Krueger et al 2023). The filtered reads were aligned to the *Tetrahymena thermophila* macronuclear genome (TGD) (Ye et al 2024) using Bowtie2 v2.3.5.1 with the parameters -- no-mixed --no-discordant (Langmead & Salzberg 2012). Only uniquely mapped reads were retained for downstream analyses. For both ChIP and Input samples, the aligned reads were sorted using Picard v2.18.29 (SortSam) and PCR duplicates were removed with MarkDuplicates (http://broadinstitute.github.io/picard/). Fragments of 100-200 bp were selected using the alignmentSieve function from deepTools v3.5.4 (Ramírez et al 2016).

Peak calling for both ChIP-seq and CUT&Tag data was performed using MACS2 v2.2.9.1 in broad mode. Peaks with a signalValue ≥ 2 and q-value ≤ 0.05 were considered reliable (Zhang et al 2008). To correct for background noise, genes lacking reliable peaks were selected as a background gene set. The average enrichment of these background genes was calculated using bamCoverage and computeMatrix from deepTools (Ramírez et al 2016). Custom Perl scripts were used to normalize gene-level enrichment based on the background signal. Heatmaps were generated using the plotHeatmap function of deepTools. Detailed enrichment information for each protein is provided in Supplementary Data 2.

### RNA sequencing and data analysis

1 × 10⁶ *Tetrahymena* cells were collected with 2 × 10⁵ *Pseudocohnilembus persalinus* cells as spike-in. RNA was extracted using the FastPure Cell/Tissue Total RNA Isolation Kit V2 (Vazyme, RC112).

RNA-seq data were preprocessed following the same procedure described above. The filtered reads were aligned to both the *T. thermophila* macronuclear genome (Ye et al 2024) and the *Pseudocohnilembus persalinus* macronuclear genome (Liu et al 2024) using HISAT2 v2.2.1 with default parameters (Kim et al 2019). Only uniquely mapped reads were retained, and the resulting alignments were sorted and deduplicated using Picard v2.18.29 (http://broadinstitute.github.io/picard/). Gene-level read counts for *T. thermophila* were quantified using featureCounts v2.0.6 (Liao et al 2014). To calculate normalization factors, the number of reads mapped to *P. persalinus* in each sample was transformed using the natural logarithm, and the arithmetic mean (n) across all samples was computed. The size factor for each sample was then determined by dividing its *P. persalinus* read count by exp(n).

Differential gene expression analysis was performed using DESeq2 (Love et al 2014), incorporating the calculated size factors. Genes with an adjusted p-value (p-adj) ≤ 0.05 were defined as significantly differentially expressed genes. Among these, genes with a log_2_(FoldChange) ≤ −1 were considered as significantly down-regulated, while genes with a log_2_(FoldChange) ≥ 1 were considered as significantly up-regulated. Volcano plots were generated using ggplot2 (Wickham 2016). Gene Ontology (GO) annotation of *T. thermophila* genes was conducted using PANNZER2, and GO enrichment analysis was carried out with TBtoolsII (Chen et al 2023). The results of the DESeq2 analysis and GO enrichment analysis are available in Supplementary Data 2.

## Data availability

All sequencing data generated in this study are available under BioProject accession number PRJNA1276626. ChIP-seq data for H2A.Z-NHA in wild-type *Tetrahymena thermophila* were obtained from Wang et al, 2025 (Wang et al 2025).

## Acknowledgement

The authors would like to thank Mr. Junhua Niu (Ocean University of China) for providing *Pseudocohnilembus persalinus* cells. Our special thanks are given to Prof. Weibo Song (Ocean University of China) for his helpful suggestions during drafting the manuscript. High-performance computing resources for data processing were provided by the Institute of Evolution and Marine Biodiversity, High-Performance Biological Supercomputing Center, and Marine Big Data Center of Institute for Advanced Ocean Study at the Ocean University of China (OUC).

## Author contribution

YW and SG conceived the study and designed the experiments; ZZ, FW and AJ performed the experiments; HL performed the bioinformatic analysis with FY; YL modified the figures with HJ; ZZ, HL and YW wrote the paper, with inputs from all authors. All authors read and approved the final manuscript.

The authors declare no conflict of interest.

## References

Allis CD, Glover CVC, Bowen JK, Gorovsky MA. 1980. Histone variants specific to the transcriptionally active, amitotically dividing macronucleus of the unicellular eucaryote, *Tetrahymena thermophila*. Cell 20: 609–17

Allis CD, Richman R, Gorovsky MA, Ziegler YS, Touchstone B, et al. 1986. hv1 is an evolutionarily conserved H2A variant that is preferentially associated with active genes. J. Biol. Chem. 261: 1941–48

Altaf M, Auger A, Monnet-Saksouk J, Brodeur J, Piquet S, et al. 2010. NuA4-dependent acetylation of nucleosomal histones H4 and H2A directly stimulates incorporation of H2A.Z by the SWR1 Complex. J. Biol. Chem. 285: 15966–77

Billon P, Côté J. 2012. Precise deposition of histone H2A.Z in chromatin for genome expression and maintenance. Biochim. Biophys. Acta Gene Regul. Mech. 1819: 290–302

Brahma S, Udugama MI, Kim J, Hada A, Bhardwaj SK, et al. 2017. INO80 exchanges H2A.Z for H2A by translocating on DNA proximal to histone dimers. Nat. Commun. 8: 15616

Bruce K, Myers FA, Mantouvalou E, Lefevre P, Greaves I, et al. 2005. The replacement histone H2A.Z in a hyperacetylated form is a feature of active genes in the chicken. Nucleic Acids Res. 33: 5633–39

Cassidy-Hanley DM. 2012. Chapter 8 - *Tetrahymena* in the Laboratory: Strain Resources, Methods for Culture, Maintenance, and Storage In Methods in Cell Biology, ed. K Collins, pp. 237–76: Academic Press

Chalker DL. 2012. Chapter 11 - Transformation and Strain Engineering of *Tetrahymena* In Methods in Cell Biology, ed. K Collins, pp. 327–45: Academic Press

Chen C, Wu Y, Li J, Wang X, Zeng Z, et al. 2023. TBtools-II: A “one for all, all for one” bioinformatics platform for biological big-data mining. Mol. Plant 16: 1733–42

Chen X, Gao S, Liu Y, Wang Y, Wang Y, Song W. 2016. Enzymatic and chemical mapping of nucleosome distribution in purified micro- and macronuclei of the ciliated model organism, *Tetrahymena thermophila*. Sci. China Life Sci. 59: 909–19

Cuddapah S, Barski A, Cui K, Schones DE, Wang Z, et al. 2009. Native chromatin preparation and Illumina/Solexa library construction. Cold Spring Harb. Protoc. 2009: pdb.prot5237

Diegmüller F, Leers J, Hake SB. 2025. The “Ins and Outs and What-Abouts” of H2A.Z: A tribute to C. David Allis. J. Biol. Chem. 301: 108154

Krueger F, Ewels P, Afyounian E, Weinstein M, Schuster-Boeckler B, et al. 2023. TrimGalore: v0.6.10 – add default decompression path (0.6.10). Zenodo.

Fujisawa T, Filippakopoulos P. 2017. Functions of bromodomain-containing proteins and their roles in homeostasis and cancer. Nat. Rev. Mol. Cell Biol. 18: 246–62

Fukuto A, Masae I, Tsuyoshi I, Jiying S, Yasunori H, et al. 2018. SUMO modification system facilitates the exchange of histone variant H2A.Z-2 at DNA damage sites. Nucleus 9: 87–94

Gao S, Xiong J, Zhang C, Berquist BR, Yang R, et al. 2013. Impaired replication elongation in *Tetrahymena* mutants deficient in histone H3 Lys 27 monomethylation. Genes Dev. 27: 1662–79

Giaimo BD, Ferrante F, Herchenröther A, Hake SB, Borggrefe T. 2019. The histone variant H2A.Z in gene regulation. Epigenetics Chromatin 12: 37

Kikuchi M, Takase S, Konuma T, Noritsugu K, Sekine S, et al. 2023. GAS41 promotes H2A.Z deposition through recognition of the N terminus of histone H3 by the YEATS domain. Proc. Natl. Acad. Sci. U.S.A. 120: e2304103120

Kim D, Paggi JM, Park C, Bennett C, Salzberg SL. 2019. Graph-based genome alignment and genotyping with HISAT2 and HISAT-genotype. Nat. Biotechnol. 37: 907–15

Kouzarides T. 2007. Chromatin modifications and their function. Cell 128: 693–705

Kreienbaum C, Paasche LW, Hake SB. 2022. H2A.Z’s ‘social’ network: functional partners of an enigmatic histone variant. Trends Biochem. Sci. 47: 909–20

Langmead B, Salzberg SL. 2012. Fast gapped-read alignment with Bowtie 2. Nat. Methods 9: 357–59

Liao Y, Smyth GK, Shi W. 2014. featureCounts: an efficient general purpose program for assigning sequence reads to genomic features. Bioinformatics 30: 923–30

Liu Y, Mochizuki K, Gorovsky MA. 2004. Histone H3 lysine 9 methylation is required for DNA elimination in developing macronuclei in *Tetrahymena*. Proc. Natl. Acad. Sci. U.S.A. 101: 1679–84

Liu Y, Niu J, Ye F, Solberg T, Lu B, et al. 2024. Dynamic DNA N^6^-adenine methylation (6mA) governs the encystment process, showcased in the unicellular eukaryote *Pseudocohnilembus persalinus*. Genome Res. 34: 256–71

Love MI, Huber W, Anders S. 2014. Moderated estimation of fold change and dispersion for RNA-seq data with DESeq2. Genome Biol. 15: 550

Luger K, Mäder AW, Richmond RK, Sargent DF, Richmond TJ. 1997. Crystal structure of the nucleosome core particle at 2.8 Å resolution. Nature 389: 251–60

Matangkasombut O, Buratowski RM, Swilling NW, Buratowski S. 2000. Bromodomain factor 1 corresponds to a missing piece of yeast TFIID. Genes Dev. 14: 951–62

Miao W, Xiong J, Bowen J, Wang W, Liu Y, et al. 2009. Microarray analyses of gene expression during the *Tetrahymena thermophila* life cycle. PLOS One 4: e4429

Mochizuki K, Novatchkova M, Loidl J. 2008. DNA double-strand breaks, but not crossovers, are required for the reorganization of meiotic nuclei in *Tetrahymena*. J. Cell Sci. 121: 2148–58

Obri A, Ouararhni K, Papin C, Diebold M-L, Padmanabhan K, et al. 2014. ANP32E is a histone chaperone that removes H2A.Z from chromatin. Nature 505: 648–53

Qiao Y, Cheng T, Zhang J, Alfarraj SA, Tian M, et al. 2022. Identification and utilization of a mutated 60S ribosomal subunit coding gene as an effective and cost-efficient selection marker for *Tetrahymena* genetic manipulation. Int. J. Biol. Macromol. 204: 1–8

Ramírez F, Ryan DP, Grüning B, Bhardwaj V, Kilpert F, et al. 2016. deepTools2: a next generation web server for deep-sequencing data analysis. Nucleic Acids Res. 44: W160–W65

Saettone A, Garg J, Lambert J-P, Nabeel-Shah S, Ponce M, et al. 2018. The bromodomain-containing protein Ibd1 links multiple chromatin-related protein complexes to highly expressed genes in *Tetrahymena thermophila*. Epigenetics Chromatin 11: 10

Strahl BD, Allis CD. 2000. The language of covalent histone modifications. Nature 403: 41–45

Surface Lauren E, Fields Paul A, Subramanian V, Behmer R, Udeshi N, et al. 2016. H2A.Z.1 Monoubiquitylation antagonizes BRD2 to maintain poised chromatin in ESCs. Cell Rep. 14: 1142–55

Tatsumi K, Sakashita G, Nariai Y, Okazaki K, Kato H, et al. 2017. G196 epitope tag system: a novel monoclonal antibody, G196, recognizes the small, soluble peptide DLVPR with high affinity. Sci. Rep. 7: 43480

Teo G, Liu G, Zhang J, Nesvizhskii AI, Gingras A-C, Choi H. 2014. SAINTexpress: Improvements and additional features in Significance Analysis of INTeractome software. J. Proteomics 100: 37–43

Tian M, Loidl J. 2018. A chromatin-associated protein required for inducing and limiting meiotic DNA double-strand break formation. Nucleic Acids Res. 46: 11822–34

Wahab S, Saettone A, Nabeel-Shah S, Dannah N, Fillingham J. 2020. Exploring the histone acetylation cycle in the protozoan model *Tetrahymena thermophila*. Front. Cell Dev. Biol. 8: 509

Wang Y, Chen X, Sheng Y, Liu Y, Gao S. 2017. N^6^-adenine DNA methylation is associated with the linker DNA of H2A.Z-containing well-positioned nucleosomes in Pol II-transcribed genes in *Tetrahymena*. Nucleic Acids Res. 45: 11594–606

Wang Y, Nan B, Ye F, Zhang Z, Yang W, et al. 2025. Dual modes of DNA N^6^-methyladenine maintenance by distinct methyltransferase complexes. Proc. Natl. Acad. Sci. U.S.A. 122: e2413037121

Wang Y, Sheng Y, Liu Y, Zhang W, Cheng T, et al. 2019. A distinct class of eukaryotic MT-A70 methyltransferases maintain symmetric DNA N^6^-adenine methylation at the ApT dinucleotides as an epigenetic mark associated with transcription. Nucleic Acids Res. 47: 11771–89

Wickham H. 2016. ggplot2: elegant graphics for data analysis. Springer-Verlag New York.

Wong LH, Tremethick DJ. 2025. Multifunctional histone variants in genome function. Nat. Rev. Genet. 26: 82–104

Xiong J, Yuan D, Fillingham JS, Garg J, Lu X, et al. 2011. Gene network landscape of the ciliate *Tetrahymena thermophila*. PLOS One 6: e20124

Ye F, Chen X, Li Y, Ju A, Sheng Y, et al. 2024. Comprehensive genome annotation of the model ciliate *Tetrahymena thermophila* by in-depth epigenetic and transcriptomic profiling. Nucleic Acids Res. 53

Zhang Y, Liu T, Meyer CA, Eeckhoute J, Johnson DS, et al. 2008. Model-based Analysis of ChIP-Seq (MACS). Genome Biol. 9: R137

Zhang Z, Ju AL, Wang YY, Jiang HZ, Liu YQ, Gao S. 2025. Bromodomain-containing proteins in the unicellular eukaryote *Tetrahymena thermophila*. Zool. Res. 46: 538–50

